# IsoformMapper: A Web Application for Protein-Level Comparison of Splice Variants through Structural Community Analysis

**DOI:** 10.1101/2025.03.05.641708

**Authors:** Alexander Vergara, Tamara Hernández-Verdeja, Pedro Ojeda-May, Leonor Ramirez, Daniel Edler, Martin Rosvall, Åsa Strand

## Abstract

Alternative splicing (AS) plays a key role in numerous cellular processes by enabling the existence of different protein isoforms with potentially different functions and fine-tuning isoform abundance. However, despite its biological importance, tools to compare the domain composition of AS-derived protein isoforms have lagged as they require both structural data and an appropriate methodology for isoform comparison. Recent advances in AI-driven protein structure prediction, particularly AlphaFold2, have made accurate structural determination of splicing isoforms easily accessible, paving the way for new tools to compare their structures and asses the functional consequences of AS at the protein structure level. Here, we present IsoformMapper, a web resource that enables researchers to study alternative splicing at the protein structural level using protein network community analysis, which captures 3D physical interactions between protein regions often missed by traditional domain analysis, allowing comparisons of isoforms across any biological system. As a proof of concept, we used our tool to a case study, in which we studied how GENOMES UNCOUPLED1 (GUN1) dependent retrograde signaling regulates plant de-etiolation through alternative splicing in Arabidopsis. In response to light, we demonstrate that GUN1 regulates the expression of specific spliceosome components, consequently modifying alternative splicing of key genes encoding components essential for establishing of photosynthesis. Additionally, the *gun1* mutant showed altered splice variant ratios for *PNSL2, CHAOS,* and *SIG5*. Our tool revealed that these isoforms have significantly different protein community structures, illustrating the impact of AS on protein function and demonstrating the practical value of the tool. (247 words)

Graphical Abstract.
IsoformMapper enables structural comparison of protein isoforms through a multi-step analysis pipeline. Starting with AlphaFold2 protein structure predictions of splice variants, the method generates weighted network representations for each isoform. The web application processes multiple AlphaFold predictions per isoform, with the conformational variability between predictions represented as weighted edges in the network representation. These networks are then analyzed using the InfoMap community detection algorithm to identify structural communities, ultimately allowing visualization and comparison of protein isoform structures through interactive network displays and alluvial diagrams. The method is applicable to protein isoforms from any biological system, providing comprehensive analysis of alternative splicing consequences at the protein level by examining structural domains as integrated units. This holistic approach enables users to evaluate potential functional differences between isoforms based on their structural organization.

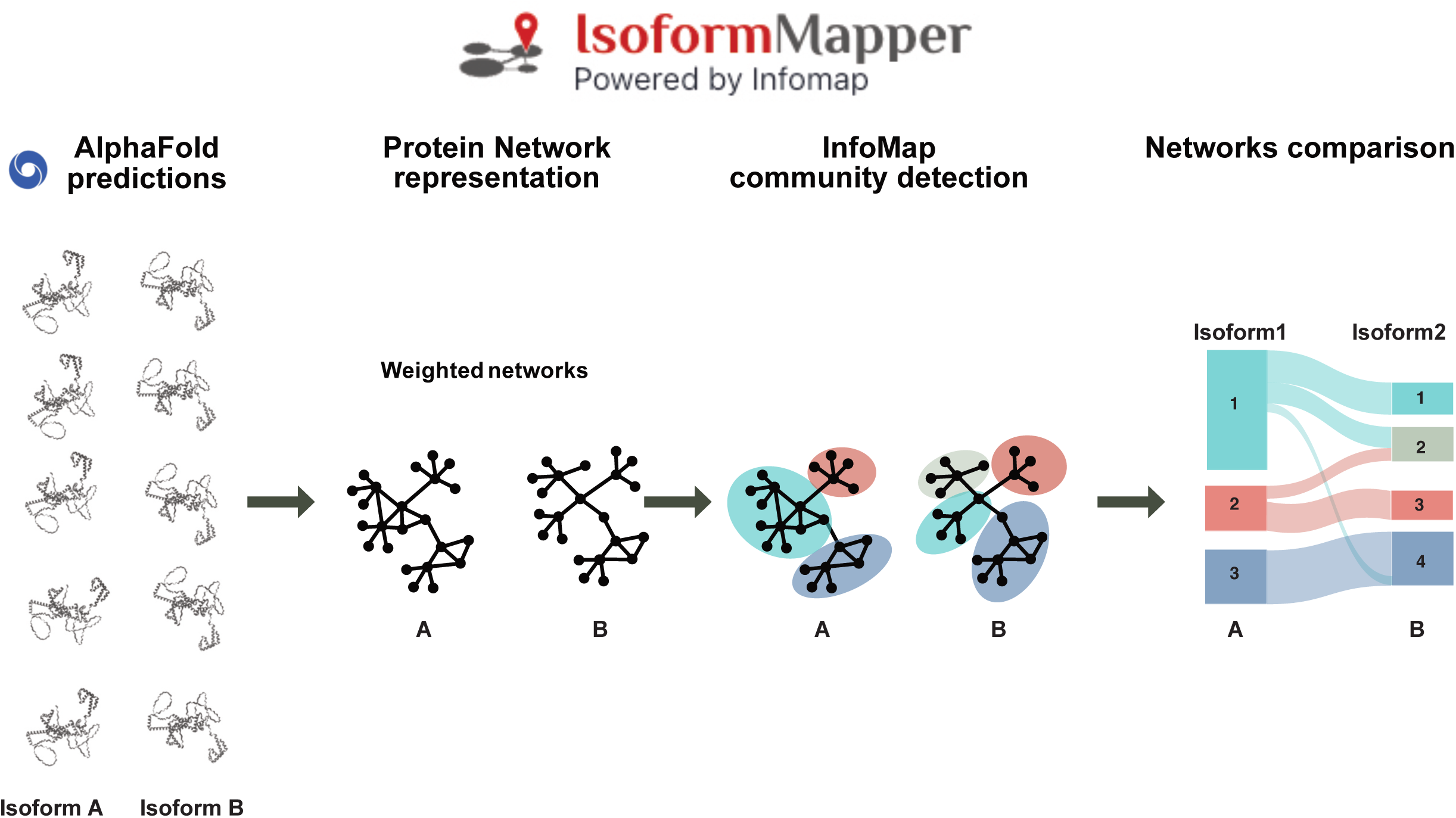

## Introduction

Alternative splicing (AS) is a post-transcriptional process that enables a single gene to generate more than one messenger RNA (mRNA) from pre-mRNA by selectively joining different sets of exons, providing a mean to increase protein diversity. Additionally, AS also regulates gene expression by introducing premature stop codons or altering untranslated regions, resulting in the degradation of unproductive transcripts or changes in mRNA stability, or detaining transcripts with retained introns in the nucleus preventing translation (Göhring *et al*., 2014; Zhou *et al*., 2024). Thus, AS can modulate mRNA levels, translation, protein activity, protein-protein interactions, affect protein post-translational modifications and subcellular localization (Reddy *et al*., 2013; Calixto *et al*., 2018; Martín *et al*., 2021; Martín *et al*., 2023; Dikaya *et al*., 2021; Kelemen *et al*., 2013; Staiger and Brown, 2013). Despite the biological importance of AS, its implications for protein structure and function remain underexplored. Addressing this gap requires accurate structural models of splicing isoforms and robust methods to analyze their structural differences and similarities.

The emergence of AlphaFold2 (Jumper *et al*., 2021) has revolutionized protein structure prediction, allowing researchers to predict multiple reliable structures for the same isoform and analyze their variability and distribution. While current bioinformatic tools can be adapted to compare protein splicing isoforms, most focus on general protein structure and function. These tools analyze primary sequences to identify domains (Blum *et al*., 2025; Sigrist *et al*., 2013), examine short segments of 3D structural motifs (Laskowski, 2022; Shi *et al*., 2007), or align 3D structures completely (Holm, 2020; Kawabata, 2003; Krissinel and Henrick, 2004). However, a comprehensive comparison of splicing isoforms requires going beyond these general-purpose sequence or structure comparisons to dissect the isoforms into domains or communities, allowing researchers to identify which domains are affected or modified between isoforms.

Analyzing structural domains in AS isoforms presents unique challenges because AS can disrupt, reconstitute, or form new domains by spatially rearranging non-consecutive residues. These changes can shift the local folding environment, potentially altering domain accessibility and stability even when the primary sequence remains mostly the same. The presence of conserved domains between isoforms suggests functional importance, while isoform-specific motifs may confer unique properties, highlighting the importance of analyzing the relationship between alternative splicing and protein structure dynamics. Comparing isoforms on a global scale is crucial to understanding how these structural differences influence function. Inconsistent residue numbering across isoforms and the challenge of handling sequence discontinuities while maintaining sensitivity to three-dimensional spatial relationships call for new methods to compare isoforms at the structural level.

To overcome these challenges, we have developed a method that compares splicing isoforms at the protein level through network analysis. Our method uses community detection analysis to define domains and subdomains based on 3D residue interactions, even when residues are not adjacent in the protein’s primary sequence. With network edge weights from multiple structure predictions, the approach can analyze different conformations. By comparing the communities associated with isoforms, we uncover large-scale changes in the interaction patterns and their functional implications.

Arabidopsis, a dicotyledonous model plant, provides an ideal system for studying the biological significance of these structural changes. When Arabidopsis seedlings germinate and emerge from the soil in darkness, they develop elongated hypocotyls with folded non-photosynthetic cotyledons (embryonic leaves). At the cellular level, these dark-grown tissues contain specialized plastids called etioplasts, which serve as precursors to photosynthetically functional chloroplasts. Here, we investigated the crucial transcriptional and post-transcriptional regulatory mechanisms that prevent premature chloroplast development during growth in the dark. These regulatory networks are essential for seedling survival, as they prevent the potentially harmful accumulation of photosynthetic components, particularly protochlorophyllide (a chlorophyll precursor) and light-harvesting proteins, which could cause photo-oxidative damage upon initial light exposure (Armarego-Marriott *et al*., 2019; Hernández-Verdeja *et al*., 2022; Kusnetsov *et al*., 2020).

In response to light, massive transcriptional changes take place, affecting nuclear and plastid-encoded genes required for chloroplast biogenesis (Jiao *et al*., 2007; Wu 2014). For establishing proper photosynthetic activity, plastid-to-nucleus (retrograde) communication coordinates nuclear and plastid gene expression (Pogson *et al*., 2008; Loudya *et al*., 2024). In the first phase of chloroplast development, photoreceptors trigger expression of nuclear-encoded genes associated with photosynthesis, known as Photosynthesis-Associated Nuclear Genes (PhANGs) (Griffin *et al*., 2022). The second phase produces fully functional photosynthetic chloroplasts and depends on a retrograde signal from the developing plastids (Dubreuil *et al*., 2018; Loudya *et al*., 2024). The timing of the second phase is linked to the activation of transcription of the photosynthesis genes encoded in the plastid genome (Diaz *et al*., 2018). Plastid transcription relies on two different RNA polymerase complexes: Nuclear-Encoded Polymerase (NEP), encoded in the nucleus, and Plastid-Encoded Polymerase (PEP), whose core subunits are encoded by the plastid genome while its DNA-binding sigma factors (SIG1-6) and other associated proteins are encoded in the nucleus (Börner *et al*., 2015). Chloroplast development likely involves a shift from NEP to PEP as the primary RNA polymerase, though the mechanisms governing this switch remain poorly understood. In green leaves, over 80% of primary plastid transcripts are transcribed by PEP (Zhelyazkova *et al*., 2012).

AS plays a critical role during plant development and acclimation to several environmental conditions such as changes in temperature and light (Reddy *et al*., 2013; Calixto *et al*., 2018; Carrasco-Lopez, *et al*., 2017; Capovilla *et al*., 2018; Dikaya *et al*., 2021; Huertas *et al*., 2019; Martín *et al*., 2021; Staiger and Brown, 2013). During photomorphogenesis, AS has been identified as an important mechanism for transcriptome diversification (Hartmann *et al*., 2016). For example, Huang et al. recently identified more than 30,000 splicing variants in etiolated seedlings exposed to white light for four hours (Huang *et al*., 2022). Despite these insights, only a limited number of studies have explored the functional consequences of specific splicing isoforms and their impact on photomorphogenesis and photosynthetic establishment (Kathare *et al*., 2021, 2022; Li *et al*., 2017; Penfield *et al*., 2010; Zhou *et al*., 1998). Phytochromes (PHYs) are photoreceptors detecting light signals that initiate signalling pathways associated with photomorphogenesis. PHYs have been reported to act upstream of AS (Shikata *et al*., 2014). However, changes to AS that are independent of photoreceptors have been identified following exposure to alternating light/dark conditions, indicating there are other light-dependent signalling pathways regulating AS. A plastid retrograde signal, that responds to light and daily light-dark cycles, was found to modulate AS of specific genes (Petrillo *et al*., 2014). Furthermore, a recent study evaluated the impact of Norflurazon, a plant herbicide known to activate retrograde signalling during chloroplast biogenesis, on AS, and discovered that AS mimics the transcriptional reactions induced by retrograde signals, while light exerts an antagonistic influence on AS (Martín, 2023).

As a proof of concept, we used our tool to analyze splicing isoforms altered in GUN1 (GENOMES UNCOUPLED1) mutant plants. GUN1 is a plastid-localized protein with a central role in retrograde signaling during chloroplast biogenesis (Hernández-Verdeja *et al*., 2018, 2020; Wu & Bock, 2021; Loudya *et al*., 2024). A model in which GUN1 signal regulates nuclear gene expression acting as a safeguard during dark-to-light transition to protect the cells from oxidative damage has been presented (Hernández-Verdeja *et al*., 2022). Our research focuses on AS within the context of chloroplast biogenesis during photomorphogenesis and GUN1-dependent retrograde signaling. We investigated how GUN1 regulates nuclear AS and how it affects transcript isoform abundance. Specifically, we analyzed stable transcript isoforms capable of encoding proteins and analyzed the effects of light exposure on these protein isoforms and their structural differences. Our results show that GUN1 regulates AS by controlling the expression of several components of the spliceosome complex, and consequently the proportion of transcript isoforms encoding protein variants associated with photosynthesis. Using our tool, we showed that these isoforms lead to significantly different protein structures, highlighting its practical utility in assessing the effects of AS on protein structure and function. Finally, we showed that GUN1 not only represses transcription of photosynthesis components in the dark and during early light response but also regulates AS to safeguard the greening process.

## Results

### IsoformMapper: A Web App for comparing protein isoforms through network structural analysis

In this study, we developed a web application that enables structural comparison of alternative splicing isoforms through analysis of their three-dimensional structures. The application generates alluvial diagrams to visualize and highlight key structural differences between isoform pairs. Figure 1 shows the IsoformMapper web app workflow **(Fig. 1)**. To run the web app available on https://www.mapequation.org/isoformmapper/:

**Figure 1.**
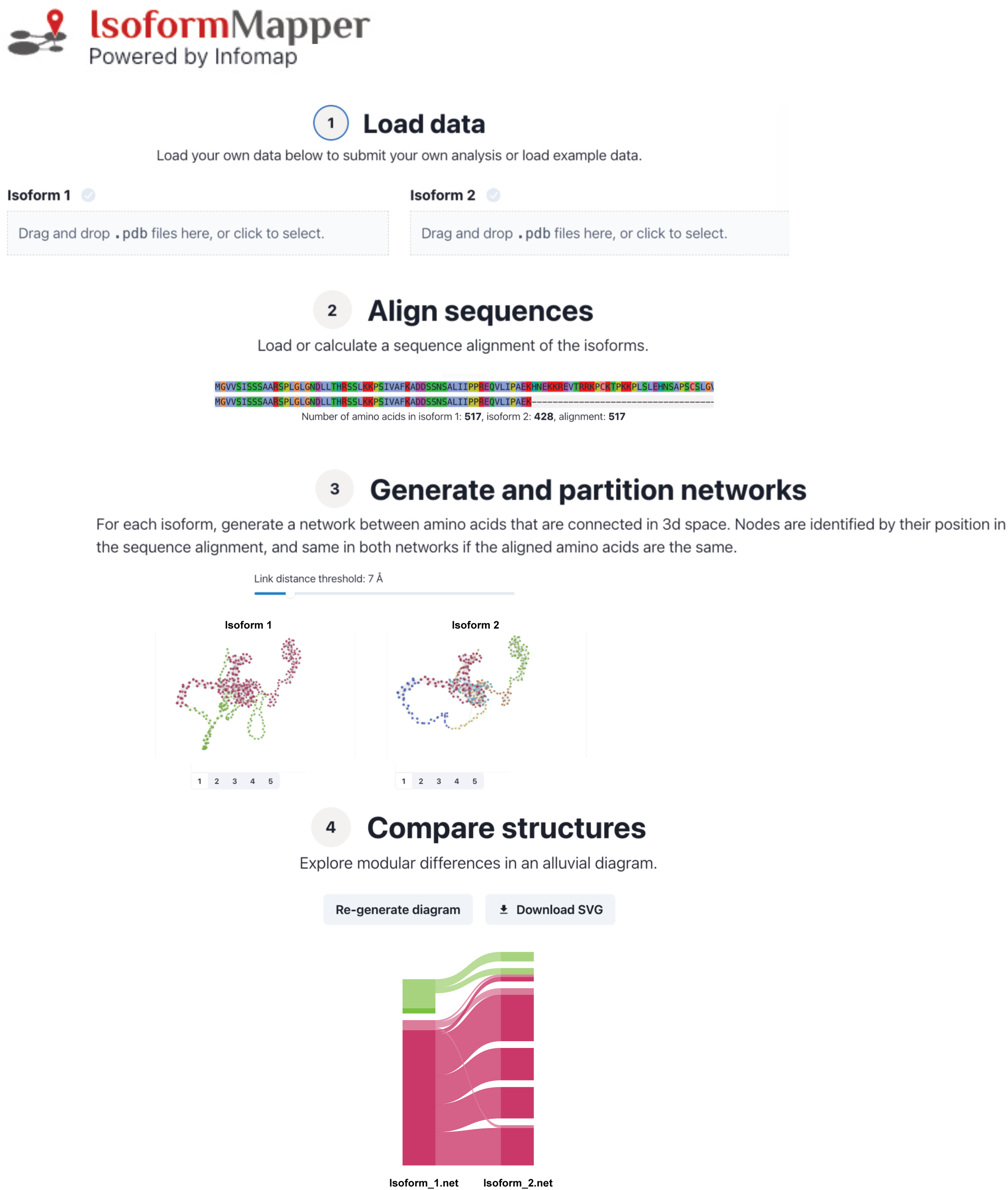
Overview of the IsoformMapper web application workflow. The application follows a five-step process for comparing protein isoform structures. In Step 1 (“Load Data”), users can upload multiple PDB files representing different conformations for each isoform to be compared. Users can also explore pre-loaded transcript pairs from our proof-of-concept study analyzing isoform switches during de-etiolation. Step 2 (“Align Sequences”) performs sequence alignment between the isoforms. Step 3 (“Generate and Partition Networks”) constructs protein networks and enables users to run InfoMap with customizable parameters to identify communities within each protein network. Users can adjust the minimum distance between amino acids alpha carbons to define network links and export the resulting networks. The application generates protein network visualizations with tabs corresponding to each uploaded PDB file per isoform, requiring users to execute InfoMap for each isoform independently. Finally, Step 4 (“Compare Structures”) visualizes the comparative analysis results through an alluvial diagram, which can be exported as an SVG file.

#### 1. Load protein 3D structures

For each isoform, drag and drop one or multiple corresponding PDB files that were obtained from AlphaFold2 to predict the 3D structure of the protein.

#### 2. Align the sequences

Click ‘align’ to generate a sequence alignment that can be visualized in its entirety using the scrollbar.

#### 3. Generate and partition networks

Click ‘Run Infomap’ to generate a network partition for each isoform. You can change the link distance threshold (default is 7 Å) and get immediate visual feedback on the 3d network.

#### 4. Generate an alluvial diagram

Click ‘Generate diagram’ to show the structural differences between the two isoforms on a modular level.

The AlphaFold2 files include per-residue confidence scores (pLDDT) that give users an indication of which parts of the predicted structure are reliable (Guo *et al*., 2022). We highlight low-confidence structures (pLDDT < 70) in the alluvial diagram using paler colors. Users can download both the underlying network files and the generated alluvial diagram as SVG files through the web interface for downstream analysis and visualization. For exploration, we have provided some examples, including the data analyzed in the manuscript.

The web app generates community partitions in weighted amino acid networks via Infomap and displays structural differences through an alluvial diagram (Holmgren *et al*., 2023; Rosvall & Bergstrom, 2010), where the N-terminal is at the top and the C-terminal is at the bottom for each isoform.

### Case study: Analyzing splicing isoform alterations mediated by GUN1 retrograde signaling during de-etiolation

GUN1 is a plastid-localized protein critical for retrograde signaling during chloroplast development. In this study, we investigated transcriptomic changes during de-etiolation by reanalyzing RNA-Seq data from wild-type and *gun1-102* mutant plants (Hernández-Verdeja *et al*., 2022). Samples included 5-day-old etiolated seedlings (0h) and seedlings that were subsequently exposed to continuous white light for 3 and 24 hours (3h and 24h, respectively). We initially compared the effects of light on wild-type seedlings. Subsequently, we contrasted wild-type plants with GUN1 mutants. For each comparison, we characterized the alternative splicing mechanisms underlying the observed transcript changes, enabling a comprehensive examination of de-etiolation and GUN1’s impact on gene expression and alternative splicing.

### Effects of de-etiolation on the transcriptome

To reveal transcriptome changes during photomorphogenesis in wild-type at both gene and transcript level, we compared by pairwise comparisons RNA-Seq data of wild-type plants. For transcripts analysis, we used the *Arabidopsis thaliana* Reference Transcript Dataset 2 (AtRTD2-QUASI) (Zhang *et al*., 2017), that enables quantification of 82,621 transcripts expression levels.

As expected, the data showed that light induction of photomorphogenesis results in massive gene expression changes, with many genes being induced or repressed by light and that these numbers were higher after 24h of light, with 14,749 differential expressed genes (DEGs) (Adjusted Pvalue < 0.05) if we consider 0h vs 3h and 0h vs 24h **(S1 Fig.)** (Jiao *et al*., 2007). To reveal the impact of alternative splicing during de-etiolation, we performed transcript isoform switch analysis by pairwise comparisons and obtained pairs of transcripts that significantly change between two time points during de-etiolation (0h/3h, 3h/24h and 0h/24h). Then, we identified the unique genes associated to those transcripts (differentially spliced genes, DSGs) **(Fig. 2a, S1 Data)**. We observed that the number of transcripts undergoing AS increased with light exposure time **(Fig. 2a)**. When we analyzed the different types of alternative splicing associated to the transcripts undergoing isoform switch, we found that intron retention (IR) was the most prevalent, followed by Alternative 3’splice site (A3) and Alternative 5’ splice site (A5) for the comparisons of all three time points **(Fig. 2b)**. We analyzed the intersections between the different lists of DSGs, and we found that the largest group with isoform switch was 0h vs 24h-specific (455 genes) **(Fig. 2c)**. Then, the second largest group corresponded to genes common between 3h vs 24h and 0h vs 24h (288 genes) while the third largest group corresponded to DSG only present in the group 3h vs 24h (274 genes), followed by 176 DSGs 0h vs 3h-specific, and only 23 DSGs were common to the three comparisons analyzed. Gene Ontology (GO) enrichment analysis for the DSGs **(Fig. 3)** revealed that genes that undergo AS during de-etiolation were enriched in functional tags related to “chloroplast”, and “RNA binding”, with more genes associated to similar terms among 0h vs 24h and 3h vs 24h DSGs. Interestingly, “RNA splicing” specific associated GO terms are only enriched in 0h vs 3h and 0h vs 24h DSGs, with several splicing regulators such as Ser-Arg (SR) undergoing IR or ES in the transition from etiolated to de-etiolating seedlings (e.g. *SR30*, *SR34a*, *RS31*). These results reveal a highly dynamic regulation of AS during photomorphogenesis, affecting genes related to chloroplast and RNA metabolism, with main changes in levels of splicing isoforms between early times (0h, 3h) and the last time point (24h) when chloroplast are fully developed.

**Figure 2.**
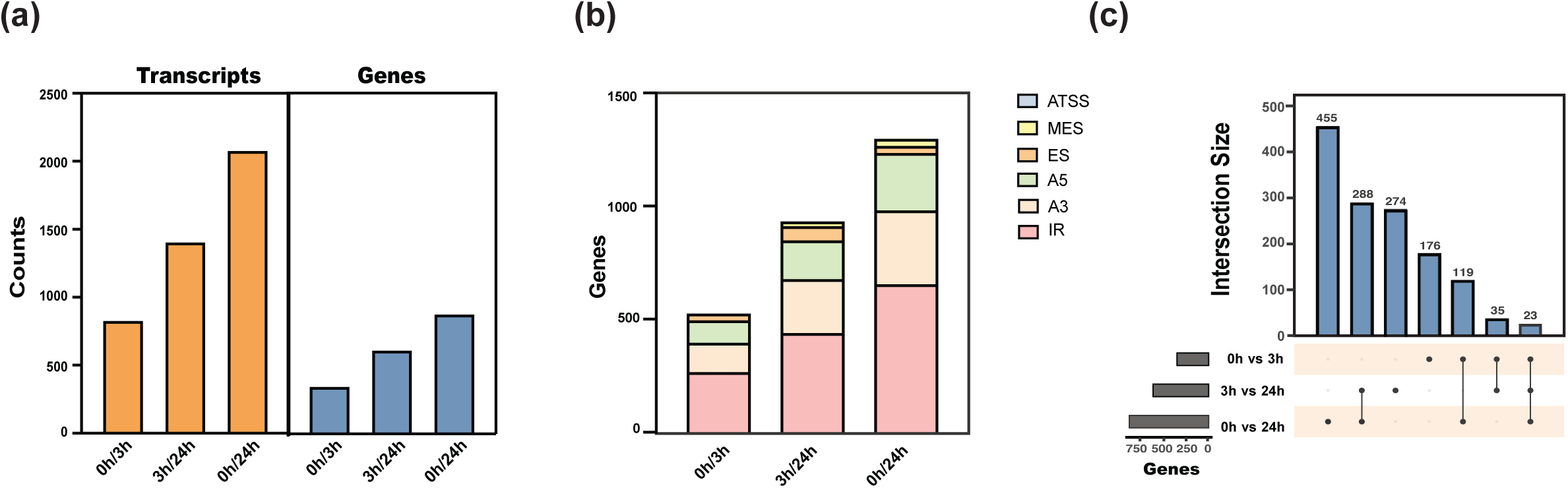
Differentially spliced transcripts and genes during de-etiolation in wild-type. (a) Differentially spliced transcripts and the respective associated unique differentially spliced genes (DSGs). (b) Statistics of DSGs and alternative splicing mechanisms. Alternative transcription start site (ATSS), Multiple exon skipping (MES), Exon skipping (ES), Alternative 5’ splice site (A5), Alternative 3’ splice site (A3), Intron retention (IR). The presence of two genes with ATTS in 0h/24h is not clearly evident in the figure due to scale of the bar plot. (c) Intersection analysis between DSGs lists. A comprehensive list of DSGs in the wild type during de-etiolation, including annotations, alternative splicing mechanisms, the corresponding transcript isoforms involved, and the respective timepoints for comparison where changes were observed, is available in S1 Data.

**Figure 3.**
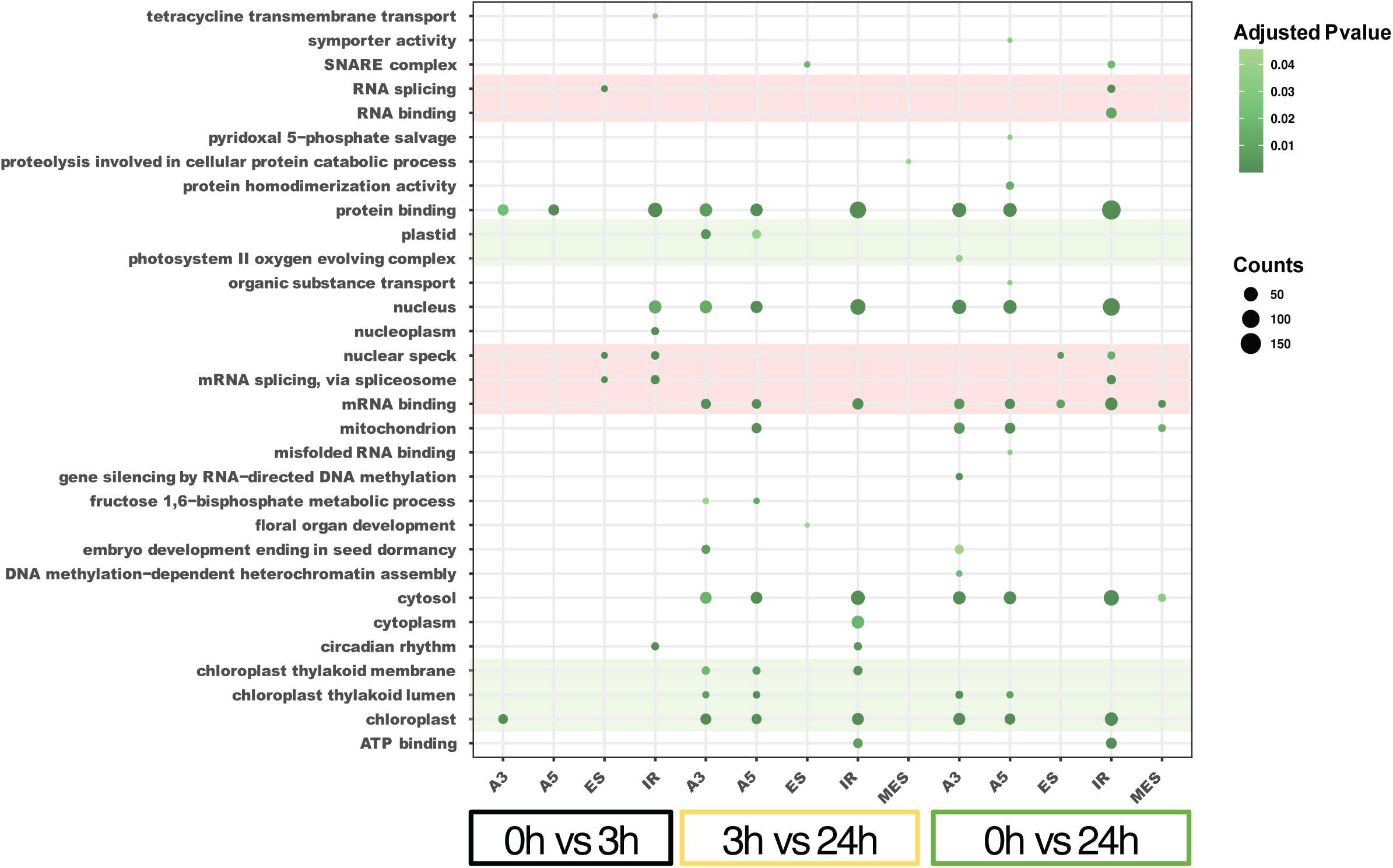
Gene Ontology enrichment analysis in differentially spliced genes (DSGs) in wild-type seedlings during de-etiolation. Lists of genes displaying differential spling events (DSGs) were grouped according to alternative splicing mechanisms and to the time of exposure to light. Alternative 3’ splice site (A3), Alternative 5’ splice site (A5), Exon skipping (ES), Intron retention (IR), Multiple exon skipping (MES). Functional tags related to photosynthesis, RNA binding and alternative splicing are highlighted in green and pink, respectively.

### GUN1 retrograde signal up-regulates expression of splicing-related genes

Our analysis revealed the regulation of AS during photomorphogenesis. Based on recent studies suggesting that a GUN1-dependent retrograde signal regulates AS (Martín G. 2023), we hypothesized that GUN1 retrograde signaling (RS) could regulate the expression of nuclear genes involved in nuclear splicing. To explore this in depth, we analyzed the previously RNA-seq data including *gun1-102* for the three time points (0h, 3h, and 24h), and performed pairwise analysis for differential gene expression between wild-type seedlings (WT) and *gun1-102* for each time point. Gene Ontology (GO) enrichment analysis of DEGs for each comparison revealed that GO terms associated with chloroplast, RNA metabolism, and AS were enriched in *gun1* down-regulated genes **(Fig. 4a)**. Interestingly, this analysis also showed that the down-regulated genes were specifically enriched in GO terms related to RNA metabolism and alternative splicing (AS) in etiolated seedlings (0h) and at 3h of light exposure, but this enrichment was not significant after 24h **(Fig. 4a)**. When we analyzed the expression of all genes with GO terms related to RNA metabolism and RNA splicing, we found that most of these genes were down-regulated in *gun1*, including genes encoding proteins that form the core of the spliceosome and are involved in AS, such as *SME1*, *SME2*, *SMD1a*, *LSM2*, *LSM7*, and some alternative splicing factors, such as *SKIP*, *PRP4KA*, *SR34* (Carrasco-Lopez *et al*., 2017; Elvira-Matelot *et al*., 2016; Capovilla *et al*., 2018; Huertas *et al*., 2019, Kanno *et al*., 2018, Wang *et al*., 2012) **(Fig. 4b)**. The results indicate that GUN1 regulates the expression of genes encoding components of the spliceosome, and suggest GUN1 has significant downstream effects on the balance between different splicing isoforms during de-etiolation and chloroplast biogenesis.

**Figure 4.**
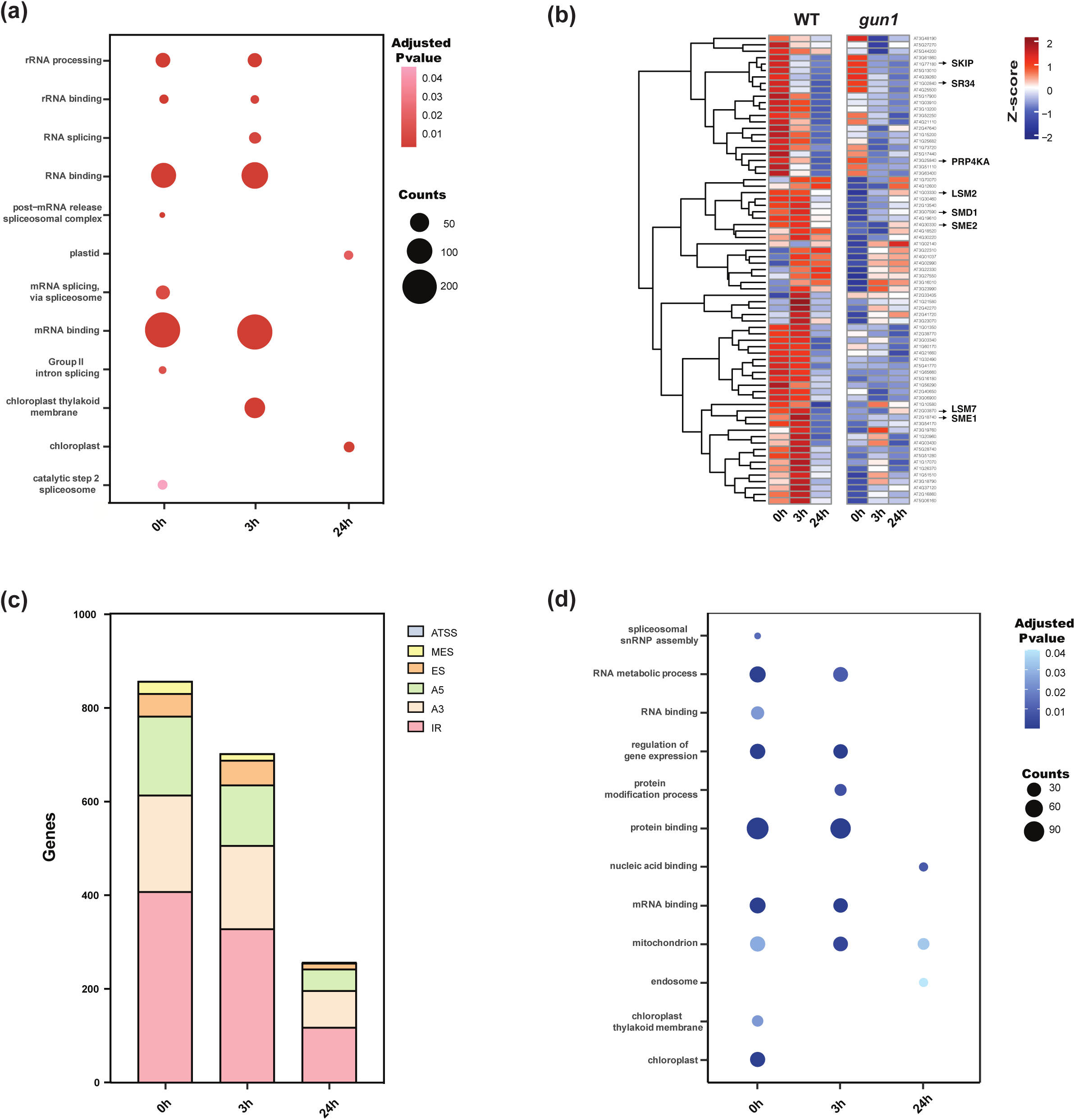
Splicing is affected in *gun-1* during de-etiolation. (a) Selected Gene Ontology (GO) terms that are enriched in *gun1* down-regulated genes. (b) Heatmap of genes involved in alternative splicing. (c) Statistics of differentially spliced genes (DSGs) between WT and *gun1* at different times of light exposure and respective alternative splicing mechanisms. Alternative transcription start site (ATSS), Multiple exon skipping (MES), Exon skipping (ES), Alternative 5’ splice site (A5), Alternative 3’ splice site (A3), Intron retention (IR). (d) Selected GO terms enriched in DSGs between WT and *gun1*. A comprehensive list of DSGs changing between wild type and *gun1* during de-etiolation, including annotations, alternative splicing mechanisms, the corresponding transcript isoforms involved, and the respective timepoints for comparison where changes were observed, is available in S2 Data.

### GUN1 affects alternative splicing during chloroplast biogenesis

Due to our observation that GUN1 regulates key spliceosomal genes at 0h and 3h, we identified differentially spliced genes (DSGs) between wild-type and *gun1-102* for the three time points: 0h, 3h, and 24h in light and examined the different AS types. We found that the highest number of AS in *gun1-102* compared to wild-type was present in etiolated seedlings (0h), 857 AS events in 572 unique genes. Subsequently, the number of these DSGs gradually decreases with exposure to light, with 703 and 258 AS events in 452 and 175 unique genes at 3h and 24h respectively **(Fig. 4c and S2 Data)**. The distribution of different AS types was similar at all time points, and as expected intron retention (IR) was the predominant AS mechanism, followed by Alternative 3’ splice site (A3) and then very closely by Alternative 5’ splice site (A5) **(Fig. 4c)**. We analyzed the enrichments of GO terms in the lists of unique genes undergoing AS for the three time points. GO terms related to RNA metabolism and alternative splicing itself were enriched, especially at 0h. Distinctly, GO terms related to chloroplast are only enriched at 0h, but not after light exposure and start of chloroplast development (3h, 24h). Analysis reveals a trend with GUN1 mainly regulating AS of genes involved in chloroplast and RNA metabolism in etiolated seedlings, functional categories that gradually lose significance with progress of light exposure and chloroplast development **(Fig. 4d)**. This role of GUN1 regulating alternative splicing is correlated with GUN1 protein levels that are higher in etiolated seedlings and decrease with chloroplast development and exposure to light until it is negligible (Hernández-Verdeja *et al*., 2022).

### GUN1 regulates the alternative splicing of light-responsive genes during de-etiolation

Since GUN1 retrograde signal is linked to chloroplast biogenesis in light, and we found differentially spliced genes (DSGs) associated with chloroplast GO annotations that changed in *gun1-102*, we decided to focus on these genes and identify DSGs within this functional category that are alternative spliced in wild-type during de-etiolation. In total, 40 genes were identified corresponding to 183 AS events. All these genes shared the following characteristics: they were nuclear-encoded genes related to chloroplast or photosynthesis, exhibit transcript changes in wild-type plants during de-etiolation, and exhibit transcript alterations in *gun1-102* plants compared to the wild-type during de-etiolation **(S3 Data)**. We also predicted premature termination codons (PTCs) and Nonsense-Mediated mRNA Decay (NMD) sensitivity of the different AS isoforms for these 40 genes **(S4 Data)**. Results show that 67% of the genes have at least one isoform with PTC, that could result in NMD, reduced stability, or altered translation efficiency. For a detailed analysis we selected five genes with AS isoforms that could be translated into proteins-without PTC-, that are directly involved in the photosynthetic process: HCEF1, PORB, PNSL2, CHAOS and SIG5. *HCEF1* encodes a chloroplast fructose 1,6-bisphosphate phosphatase; *PORB* encodes one of the protochlorophyllide oxidoreductases; *PNSL2* encodes a subunit of the photosynthetic NAD(P)H dehydrogenase complex; *CHAOS* encodes the chloroplast signal recognition particle 43; and *SIG5* encodes a sigma factor required for PEP-dependent transcription of light-dependent genes. All these five genes present pairs of transcripts that changed during de-etiolation and have different patterns in wild-type and *gun1-102* **(Fig. 5)**. We validated expression of transcript isoforms for these five photosynthesis-related genes by qPCR in WT seedlings **(S2 Fig.)**. Analysis of the different protein products associated with the transcript isoforms showed that *HCEF1* and *PORB* isoform protein products result in the same protein sequence, while *PNSL2, CHAOS* and *SIG5* isoforms produce different protein products **(S3 Fig.)**. These three genes each have at least two transcript isoforms that undergo isoform switching: PNSL2.1 and PNSL_c1, CHAOS_P1 and CHAOS_s1, and SIG_P1 and SIG_c2, respectively. The results reveal that GUN1-dependent signal regulates AS pattern of photosynthetic transcripts that undergo AS during de-etiolation.

**Figure 5.**
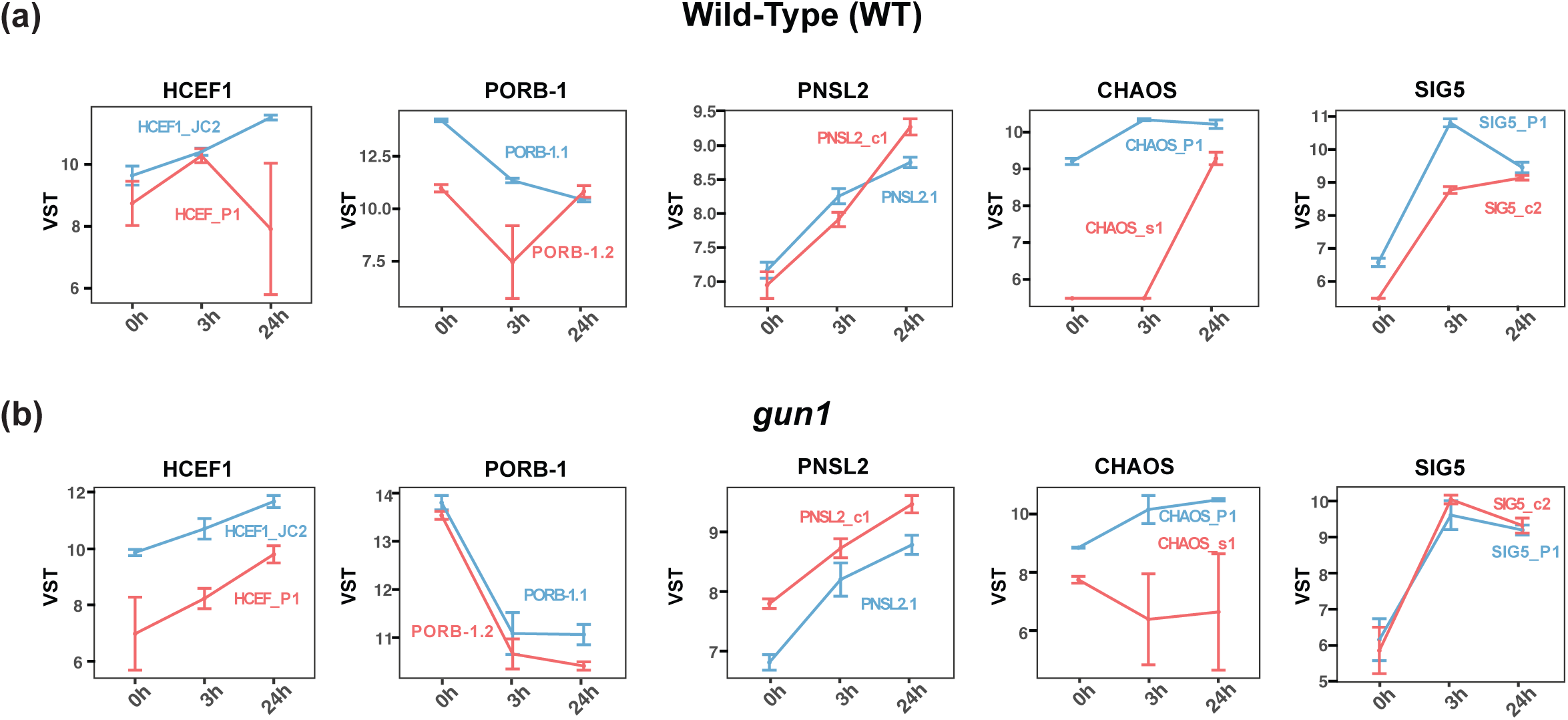
Expression profiles of photosynthesis related transcripts undergo isoform switching in wild-type and *gun1* during de-etiolation. Each plot contains transcripts identified as undergoing isoform switch. RNA-Seq normalized expression levels obtained by variance stabilizing transformation (VST) for transcripts with isoform switch between wild-type (a) and *gun1* (b) are represented. Errors bars represent SD values. Names of the respective transcripts are shown in the same color of the curve that they represent.

### Proof of Concept: Exploring Isoform Structural Changes During De-etiolation using IsoformMapper

We used our web app IsoformMapper to compare different protein isoforms encoded by three genes crucial for photosynthesis: PNSL2, CHAOS, and SIG5. These genes play key roles in the de-etiolation process and the subsequent establishment of photosynthetic machinery. We used the distance threshold of 7 Å, which was suggested in previous works as an optimal value (Barloti *et al*., 2007; da Silveira *et al*., 2009; Yan *et al*., 2014). While we have set that distance as the default in our web app, our app is capable of accommodating user-defined values. As weight of the link between two amino acids, we used the average values of distances over five different AlphaFold2 conformations. Users can generate any number of conformations, load them in IsoformMapper and the app will process these to calculate a weighted average distance. This average is then used to weight the corresponding protein networks (Yan *et al*., 2014).

For PNSL2 we identified two isoforms that could be translated to proteins, PNSL2.1 (190 aa) and PNSL1_c1 (133 aa). After 24 hours light exposure the smaller isoform PNSL1_c1 is the dominant splice variant in wild-type, while in *gun1-102* this smaller variant is the predominant form at the three time points **(Fig. 5)**. Our IsoformMapper analysis revealed that PNSL2.1 contains four main communities corresponding to distinct structural domains: communities 1 (red) and 2 (green) form alpha helices, while communities 3 (dark orange) and 4 (cyan) represent unstructured domains with high flexibility **(Fig. 6a & S4a Fig.)**. A splitting of domains 3 and 4 was observed in PNSL2.1, resulting in a different arrangement of domains 2, 3, and 4 in PNSL2_c1. This shorter PNSL2_c1 isoform lack of substantial amino acid sequences present in communities 1 and 2 of the longer PNSL2.1 isoform. Also, PNSL2_c1 contains unique amino acids in its community 1 (green-gray) that are not present in PNSL2.1. However, in addition to its specific C-terminal segment (gray), it also possesses a small region that corresponds to amino acids in community 1 of PNSL2.1, revealing a small sequence conservation between both isoforms generated by a 5′ alternative splice site (A5′) and that these amino acids participate in specific interactions within this community in the smaller isoform **(Fig. 6a**, **S4a Fig and S2 Data.)**.

**Figure 6.**
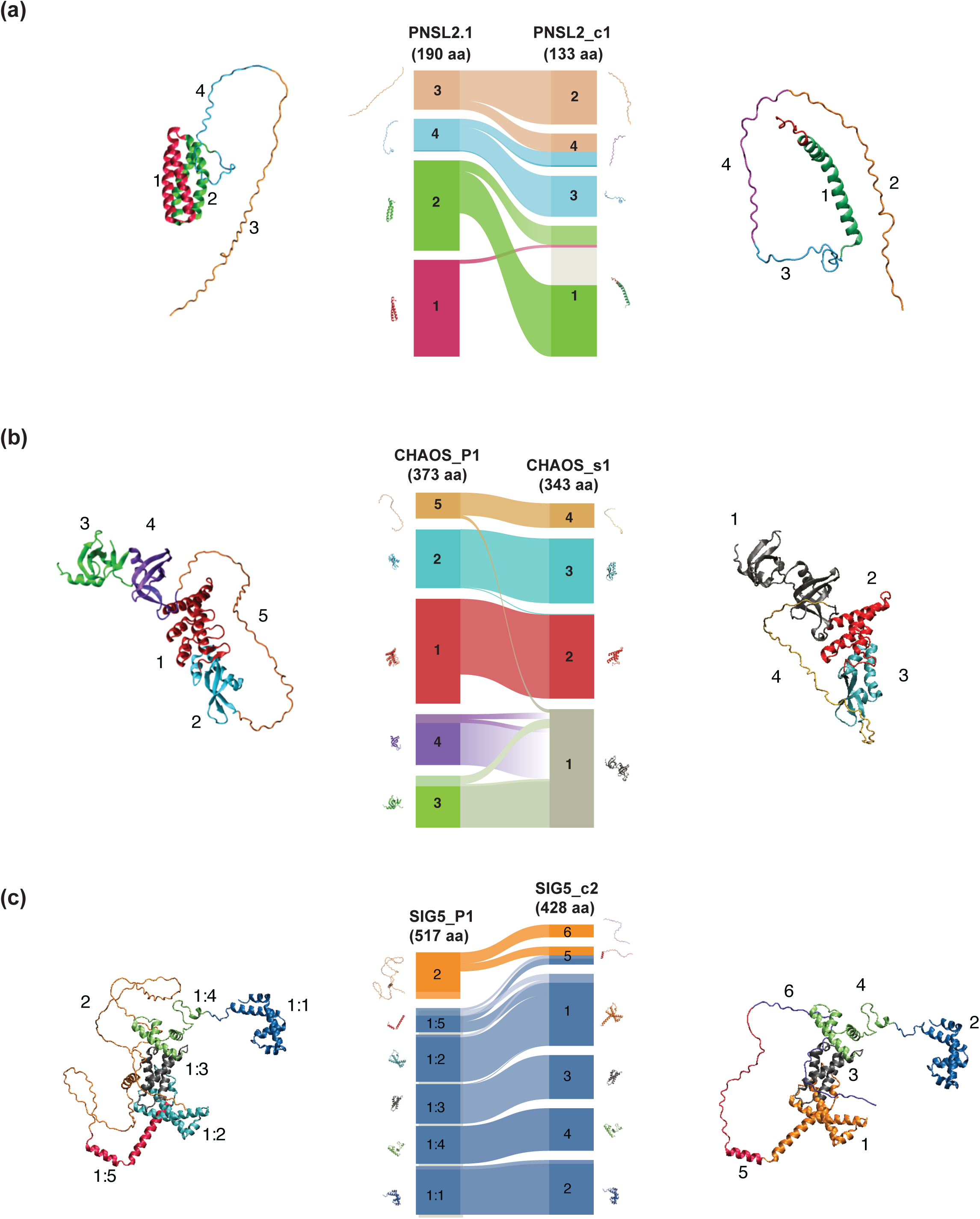
Network-based comparative analysis of protein isoforms. Protein isoforms that were identified as undergoing transcript isoform switch were analyzed by protein network analysis and structural community analysis (see methods). Network representations of different AlphaFold isoform models were obtained from amino acid networks weighting five AlphaFold model conformations. Infomap multilevel community (Rosvall & Bergstrom, 2008, 2011) and mapping change analysis (Holmgren et al., 2023; Rosvall & Bergstrom, 2010) were used to compare the networks between them using alluvial plots. In these plots, the N-terminus and C-terminus of the respective protein isoforms were represented at the top and bottom of alluvial diagrams, respectively. (a) PNSL2, (b) CHAOS and (c) SIG5.

We identified two isoforms for CHAOS, CHAOS_P1 (373 aa) and CHAOS_s1 (343 aa), where the larger isoform CHAOS_P1 is the only form present in darkness whereas the smaller splice variant CHAOS_s1 is strongly induced by light exposure, while in *gun1-102* the level of the smaller isoform is also high at 0h and 3h **(Fig. 5)**. For CHAOS IsoformMapper identified five main communities for its larger isoform CHAOS_P1 (373 aa) and four communities for the smaller isoform CHAOS_s1 (343 aa), where community 5 in CHAOS_P1 and 4 in CHAOS_s1 (beige) are unstructured domains. Community 1 of CHAOS_P1 included a region that was lacking in CHAOS_s1, and this generates structural changes between the two protein isoforms. CHAOS_s1 shapes its community 1 by the merge of communities 4 and 3 plus some few amino acids of community 5 of CHAOS_P1. Also, the analysis showed that specific amino acids from community 2 of CHAOS_P1 are repositioned in community 2 of CHAOS_s1, demonstrating their involvement in distinct interactions within this community in the shorter isoform. **(Fig. 6b)**.

We identified two isoforms for SIG5, SIG5_P1 (517 aa) and SIG5_c2 (428 aa). The larger isoform SIG5_P1 is the dominant splice variant in the dark and during early light response whereas following 24 hours equal levels of the two splice variants SIG5_P1 and SIG5_c2 were found **(Fig. 5)**. In contrast, in *gun1-102* the levels of both isoforms remain similar during de-etiolation **(Fig. 5)**. By using IsoformMapper we revealed that the larger isoform SIG5_P1 (517 aa), has two main communities. Community 1 (blue) can be split into five subdomains (1:1, 1:2,1:3,1:4 and 1:5), which all have either a beta sheet structure or an alpha helix. Community 2 (golden color) is rather unstructured and very flexible and includes amino acids that are lacking in the SIG5_c2 isoform **(S4c and S5 Fig.)**. The smaller isoform SIG5_c2 (428 aa) have six communities. Their communities 5 and 6 correspond to a split of SIG5_P1 community 2, being SIG5_c2 community 5 formed by a merge with some segment of community 1:5 of SIG5_P1 as well. The two isoforms have different distributions of communities. SIG5_c2 has a shorter, flexible segment compared to the larger SIG5_P1 isoform. This difference in structure, particularly the higher number of interactions within the unstructured segment of SIG5_P1 community 2 (golden color), influences the overall protein structure and may affect its function **(Fig. 6c)**. Taken together our analysis show that pairs of protein isoforms vary considerably at a three-dimensional level, forming distinct interactions within their amino acids, even in regions that are common between the two forms. Potentially this could result in different activities, localization or interactions that that could be modulated during de-etiolation and through GUN1 retrograde signal.

### Addressing potential consequences of the SIG5 isoform switch by molecular dynamics modelling

Since SIG5 plays an essential role in the chloroplasts by regulating the expression of plastid encoded genes associated with photosynthesis, we decided to deepen our study on the consequences of alternative splicing (AS). To address how the differences between the two SIG5 isoforms could potentially affect SIG5 function we performed a molecular dynamics analysis, in which we modeled the interaction of the two SIG5 isoforms, SIG5_P1 and SIG5_c2 with a segment of the light-responsive promoter (LRP) located upstream of the psbD gene, a well-known target of SIG5 that encodes the D2 subunit of the photosystem II reaction center complex (Kanamaru *et al*., 2004; Tsunoyama *et al*., 2004). The LRP promoter has highly conserved regulatory sequences, such as bacterial-like elements −10 and −35 and a sequence called AAG-box, which participates in the recruitment of the transcriptional machinery (see methods, Baba *et al*., 2001; Kanamaru *et al*., 2004, Thum *et al*., 2001). We used InterProScan to analyze the protein domains in both SIG5 protein isoforms and identified three DNA binding domains that are present in both protein isoforms associated with the so-called Winged helix-like DNA-binding domain superfamily (Gajiwala *et al*., 2000; Quevillon *et al*., 2005). These DNA-binding domains have the same amino acids sequences in both isoforms, but different coordinates because of their different sizes (see methods). We represented these domains by blue, red, and black colors, from the N-terminal to the C-terminal **(Fig. 7a)**. We found that the interaction profiles of the two isoforms were different even though both simulations started from similarly oriented protein-DNA structures. The results showed that the black domain in SIG5_c2 exhibited a close interaction with the DNA, along with the red and blue domains. In comparison, all three domains in SIG5_P1 demonstrated a greater distance from the DNA during the simulation time **(Fig. 7 a, b)**. This difference in the interaction with DNA may be explained by the fact that SIG5_P1 has an extra segment that adds more interaction possibilities within the segment. Interestingly, the unstructured region found by AlphaFold2 predicted conformations with a more compact conformation upon binding to the DNA. Possibly variations in the unstructured region could fine tune DNA binding affinities. Potentially, the two splice variants of SIG5 could generate proteins with different DNA binding abilities and hence alter recruitment of the PEP core complex and consequently plastid gene expression. In the *gun1-102* mutant line, SIG5_P1 and SIG5_c2 showed equal expression levels all through the de-etiolation process which contrasts with wild-type where SIG5_P1 is the dominant splice variant in the dark and during early light response **(Fig. 5)**. We compared the expression levels of chloroplast encoded genes regulated by the PEP (Ji *et al*., 2021) between wild-type and *gun1-102* using the RNA-seq data **(Fig. 7C)**. We found that most of the analyzed genes were significantly down-regulated in *gun1-102,* compared to wild-type indicating that correct balance between the two SIG5 isoforms might be important for correct gene expression in the plastids during the greening process.

**Figure 7.**
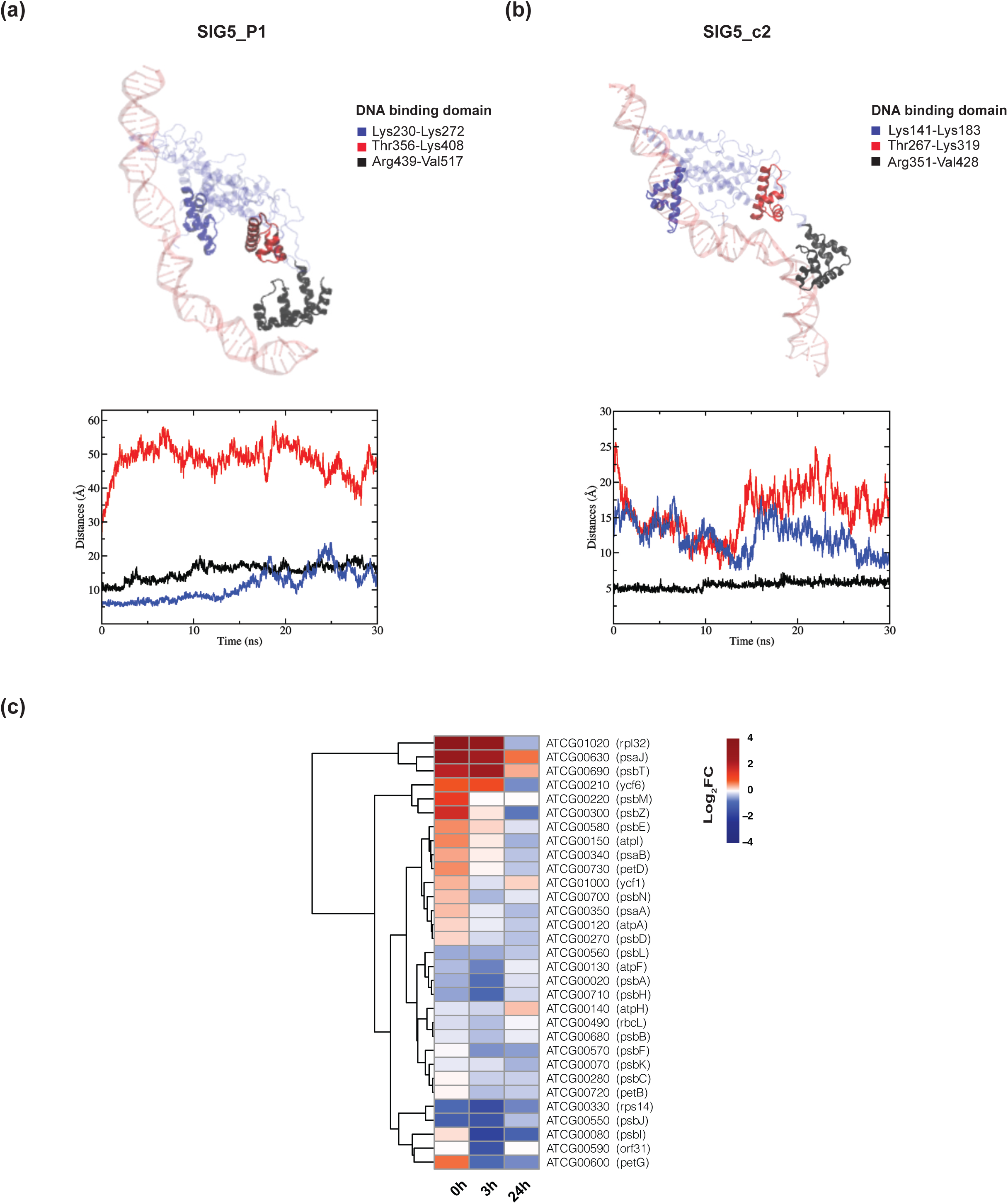
SIG5 protein isoforms and consequences of isoform switching. (a,b) Results of molecular dynamics simulations for SIG5 protein isoforms, SIG5_P1 and SIG5_c2, interacting with DNA. DNA binding domains of SIG5 present in both isoforms are highlighted in bold colors. The DNA fragment interacting with the isoforms is represented in soft red (see methods). (c) Heatmap of chloroplast-encoded genes regulated by plastid-encoded polymerase (PEP) comples, representing changes in expression between wild-type (WT) and *gun1* mutant in Log_2_FC. Up-regulated and Down-regulated genes in *gun1* compared to WT are shown in red and blue respectivelly.

## Discussion

Despite the importance of AS in generating protein diversification and its impact on isoform abundance, current methodologies are deficient in effective tools for analyzing protein isoforms at the structural level. While conventional methods, including sequence-based domain identification (Blum *et al*., 2024; Sigrist *et al*., 2013), structural motif analysis (Laskowski, 2022; Shi *et al*., 2007), and complete 3D structural alignments (Holm, 2020; Kawabata, 2003; Krissinel and Henrick, 2004), have been invaluable for general protein characterization, these methods have not been specifically designed for this purpose and often overlook the substantial structural variations that can exist between isoforms that share similar segments.

To address this gap, we have developed IsoformMapper, a web application that examines the structural implications of AS on protein isoforms through network community analysis. One limitation of the IsoformMapper resource is that it does not calculate the structural features of splicing isoforms directly, requiring users to independently determine these structures. This prerequisite step could potentially introduce variability in structural predictions depending on the methodologies employed by different users. Nevertheless, this limitation is mitigated by the fact that the web app can be fed with several structure predictions per isoform. IsoformMapper implements a comprehensive pipeline that may processes several AlphaFold2 structure predictions, constructs amino acid networks based on user-defined distance thresholds, identifies structural communities, and visualizes protein isoform differences and similarities using alluvial diagrams. By decomposing protein isoforms into functional domains and communities, our method enables precise analysis of alternatively spliced regions and their structural implications, providing researchers with new insights into the functional consequences of alternative splicing events.

Our application is freely accessible at https://www.mapequation.org/isoformmapper, enabling researchers to analyze any pair of protein isoforms by uploading their corresponding PDB files and allowing the analysis of one or more structure predictions per isoform. These structure files can be obtained through online services such as ColabFold (Mirdita et al., 2022) or through installations on institutional/computational servers. This accessibility and versatility make IsoformMapper an essential tool for investigating protein isoform diversity and its biological significance across diverse biological systems.

To demonstrate the capabilities and biological relevance of IsoformMapper, we applied our tool to investigate how GENOMES UNCOUPLED1 (GUN1) dependent retrograde signaling modulates plant de-etiolation through alternative splicing in Arabidopsis. This case study not only showcases the practical application of our method but also provides valuable insights into a fundamental biological process where alternative splicing plays a crucial regulatory role.

The AS mechanism is tightly regulated by the spliceosome, a highly conserved multi-megadalton ribonucleoprotein (RNP) complex composed by 5 ribonucleoproteins and a large number of auxiliary proteins that recognise conserved sequences of the pre-mRNA (Liu *et al*., 2022; Wahl *et al*., 2009). Both the conformation and composition of the spliceosome are highly dynamic, providing flexibility for the splicing machinery. Also, alternative splicing depends on the RNA sequences and protein regulators such as Ser-Arg (SR) proteins and heterogeneous nuclear RNPs (hnRNPs), their post-translational modifications and their combinations (Wahl *et al*., 2009). The components of the core spliceosomal RNPs, including both proteins and small-nuclear RNAs, as well as auxiliary and regulatory proteins, have been identified in plants (Dikaya *et al*., 2021; Liu *et al*., 2022; Perea-Resa *et al*., 2012).

Studies have demonstrated that environmental changes can modulate both the expression and protein levels of core and regulatory spliceosome components. For example, low temperature conditions trigger changes in the LSM nuclear complex, leading to widespread effects on both constitutive and alternative splicing patterns (Carrasco-López *et al*., 2017). Similarly, the expression of the core spliceosome component *SME1* has been shown to be altered by cold conditions, with the protein accumulating in the nucleus at low temperatures to modify the splicing landscape and regulate cold acclimation (Capovilla *et al*., 2018; Huertas *et al*., 2019). Thus, environmental conditions can modulate alternative splicing through spliceosome alterations, thereby affecting the transcriptome, protein functions, and ultimately plant cellular metabolism. In addition, the pre-mRNAs of the Arabidopsis genes that encode serine/arginine-rich splicing regulators (SR), are themselves extensively alternatively spliced, highlighting the complex regulatory nature of the splicing process. This complexity is also exemplified by the fact that 15 SR genes can produce approximately 95 different transcripts in response to hormones and temperature stress (Palusa *et al*., 2007), highlighting the critical importance of understanding both the detailed changes in AS patterns and their functional implications at the protein level.

De-etiolation, a fundamental developmental transition when seedlings emerge from darkness into light, represents one of the most dramatic environmental adaptations in plants. During the de-etiolation process we found that 2,651 transcripts corresponding to 1,096 genes, underwent an isoform switch **(Fig. 2).** Interestingly, our analysis of alternatively spliced genes revealed enrichment in genes encoding photosynthetic components. Also, the analysis of these splicing events indicates that alternative splicing plays a key role in controlling the establishment of photosynthesis during the greening process. **(Fig. 3).** The *gun1* mutant impaired in retrograde signaling displayed an AS landscape that was significantly altered compared to wild type, especially in the dark. GUN1 regulates the expression of critical and regulatory components of the spliceosome complex such as *LSM2*, *LSM7*, *SME1* and 2, *SKIP*, and *SR34*, and as a consequence, affects AS during de-etiolation and the proportion of transcript isoforms encoding proteins variants associated with photosynthesis. Thus, in addition to repressing transcription of photosynthesis components in the dark and during early light response, GUN1 regulates AS to safeguard the greening process.

We found that during the de-etiolation process, several SR proteins, including *SR30*, *SR34a*, and *RS31*, undergo intron retention (IR) or exon skipping (ES) (**Fig. 2 & S1 Data**). GUN1 modulates both the expression and alternative splicing of spliceosomal components in response to light. The primary splicing mechanism affected in the *gun1* mutant was intron retention (IR), which has been observed in other studies as the most common splicing mechanism in Arabidopsis (Dikaya *et al*., 2021; Martín *et al*., 2021; Carrasco-Lopez *et al*., 2017). We identified 40 genes associated with plastids and photosynthesis that undergo isoform switch during de-etiolation in wild-type **(S3 and S4 Data)** that also showed differential splicing in the *gun1* mutant. The genes subject to isoform switch during the greening process encode components involved in key processes during greening such as oxidative stress response, protein degradation, assembly of the photosynthetic complexes, tetrapyrrole biosynthesis and plastoquinone biosynthesis. Our web resource provides comprehensive proof-of-concept examples showcasing the changes between transcript pairs associated with all these genes, allowing users to visualize the structural effects using our methodology. To demonstrate IsoformMapper’s capabilities, we performed a detailed analysis of three genes encoding key photosynthetic components, *PNSL2*, *CHAOS*, and *SIG5* that produce different proteins isoforms. PNSL2, which is involved in the photosynthetic electron transport, has a larger protein isoform with an alpha helix domain that is completely missing in the shorter isoform resulting in a significantly different protein structure **(Fig. 6)**. Furthermore, we are making available the complete expression profiles of all Arabidopsis transcripts during de-etiolation in both wild-type and *gun1* mutant backgrounds, enabling users to examine transcripts in our transcriptomic dataset while simultaneously exploring the structural analysis results from our example cases in IsoformMapper resource (https://deetiolation-transcripts.serve.scilifelab.se/app/deetiolation-transcripts). The smaller isoform PNSL1_c1 is the dominant splice variant in light, which indicates that this alpha helix is not important for the function of PNSL2 in light and that its absence allows other segments of the protein to perform its function differently when the plant is exposed to light. The smaller splice variant of CHAOS, which is involved in chloroplast protein import, is strongly induced by light exposure. This suggest that the import of proteins for photosynthesis is more efficient when this isoform is present **(Fig. 6)**. There are 6 sigma factors in Arabidopsis (SIG1-6) and SIG5 controls the expression of at least 12% of chloroplast-encoded genes (Noordally *et al*., 2013). The SIG5 protein isoforms are flexible multidomain proteins consisting of two or more well-folded domains connected via flexible linkers. These linkers may result in large scale inter-domain motions related to the protein function and may also influence the flexibility of the protein to interact with DNA (Thomasen & Lindorff-Larsen 2022). While this study focused on analyzing static isoform structures, the application’s capabilities extend to examining conformational dynamics through molecular dynamics (MD) simulation trajectories. Molecular dynamics modelling showed that the smaller splice variant SIG5_c2 interacts closer to DNA than the larger SIG5_P1 **(Fig. 7a, b)**. The difference in DNA-protein interaction could be explained by the extra segment that adds more interaction possibilities within the SIG5 protein. Additionally, our findings indicate that SIG5_P1 interacts with DNA in a more dynamic manner, potentially stimulating induction of transcription of chloroplast-encoded genes regulated by PEP **(Fig. 7c)**. The *gun1* showed a difference in the SIG5_P1/SIG5_c2 ratio compared to wild-type with a relatively higher level of SIG5_c2. The changes in the ratio between the SIG5 isoforms could potentially give rise to the observed reduced expression of plastid encoded genes in *gun1* **(Fig. 7)** and possibly provide an explanation for the reduced greening observed in *gun1* (Susek *et al*., 1993). In summary, our model suggests that a GUN1-dependent retrograde signal from etioplasts and early developing chloroplasts regulates alternative splicing in the nucleus, affecting transcripts signatures that can affect stability, translation and if translated, protein domains and functionality **(Fig. 8)**. Thus, our results position alternative splicing downstream of GUN1 and show how alternative splicing provides an additional mechanism for coordinating the expression and protein isoforms from both the nucleus and the plastids.

**Figure 8.**
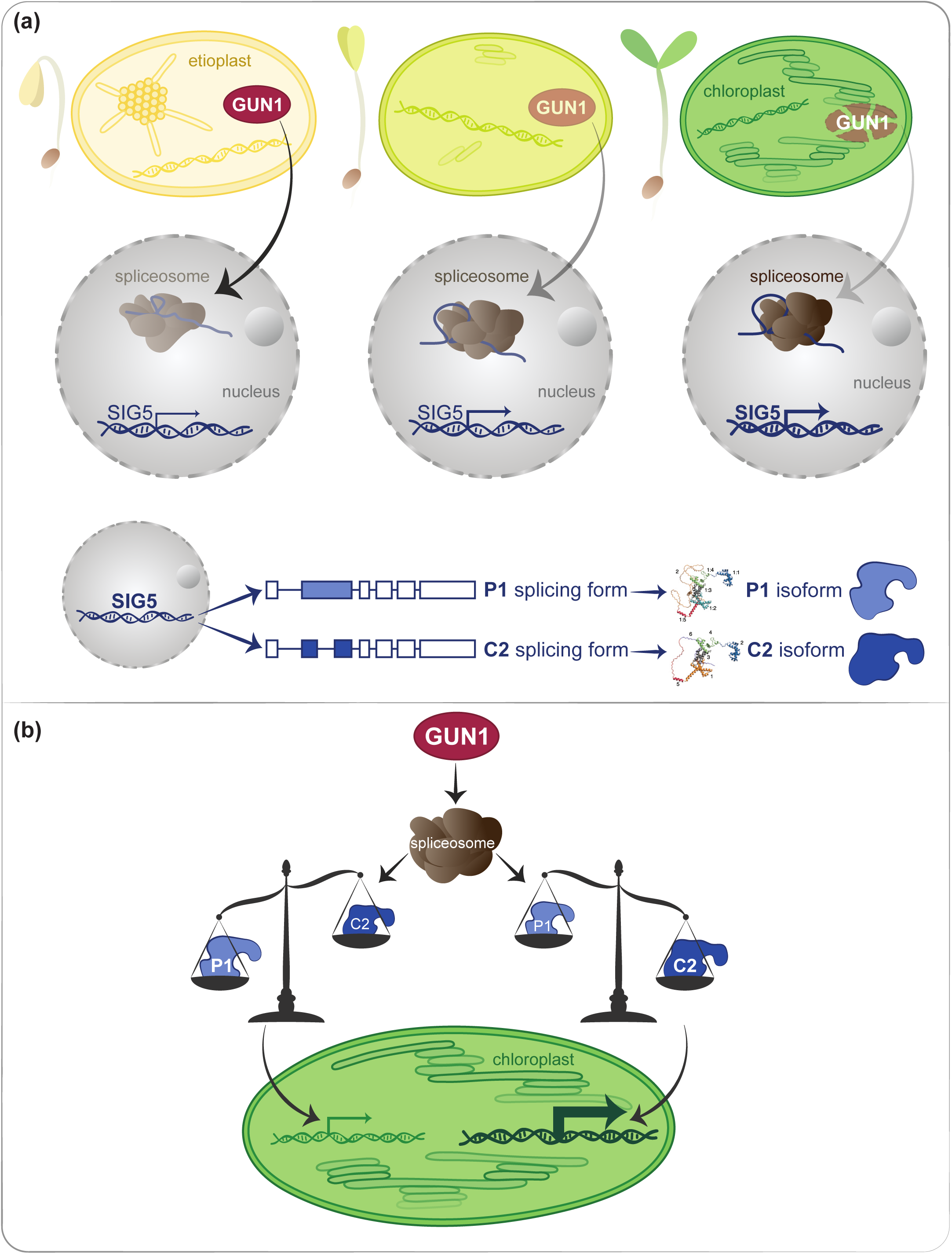
Model of GUN1 control over photosynthesis establishment. (a) Scheme representing decreasing levels of GUN1 protein with progression of light exposure and its regulation over the spliceosome composition. SIG5 is encoded in the nucleus, and levels of both isoforms increase with light exposure, coinciding with a greater induction of expression of genes encoded in the plastid. The two transcript isoforms of SIG5 undergoing isoform switch, SIG5_P1 and SIG_c2, are represented with a box model, showing differences in the N-terminus region of both isoforms. (b) The balance of both SIG5 isoforms is shown as a key element. In darkness, levels of both isoforms are low, with higher levels of SIG5_P1. After 3 hours of light exposure, levels of SIG5_P1 are much higher relative to SIG_c2, with both isoforms strongly induced by light.

## Materials and Methods

### IsoformMapper Web App development

The IsoformMapper web application, including the Infomap community detection algorithm (Rosvall & Bergstrom, 2008, 2011), runs directly in the user’s web browser using JavaScript. The web app internally aligns the respective sequences using the BioMSA JavaScript library (https://github.com/ppillot/biomsalign) and creates consensus node IDs that allow monitoring of equivalent amino acids between both sequences. After constructing the weighted amino acid networks, the web app identifies communities with Infomap and visualizes structural difference with an alluvial diagram (Holmgren *et al*., 2023; Rosvall & Bergstrom, 2010). The web app is available on https://www.mapequation.org/isoformmapper/

### RNA-Seq analysis

For the present study, RNA-seq data that was previously published (Hernández-Verdeja *et al*., 2022) was re-analyzed at genes and transcripts level and subjected to alternative splicing analysis. The raw RNA-Seq data was reanalyzed using the following methods and steps. Trimming was performed by *Trimmomatic* (v.0.39) (Bolger *et al*., 2014). Quality control was performed with *FastQC* (v0.11.4) (Andrews 2010). *SortMeRNA* (v2.1) was employed for filter RNA contaminants (Kopylova *et al*., 2012). To obtain differential expressed genes, RNA-Seq reads were mapped to Araport11 reference genome (Cheng *et al*., 2017). To obtain transcript-level quantification RNA-Seq reads were aligned against Arabidopsis Thaliana Reference Transcript Dataset 2 (AtRTD2-QUASI) (Zhang *et al*., 2017). Counts of transcripts quantification were obtained with *Salmon* (v.1.6.0) (Patro *et al*., 2017). Transcript expression levels were summarized at gene level using *tximport* (v.1.22.0) Biocoductor package (Soneson *et al*., 2015). To obtain normalized gene expression levels by variance stabilizing transformation (VST) and to perform differential gene expression analysis, *DESEQ2* R (v.1.34.0) library was used (Love *et al*., 2014). Transcript model visualizations were graphed with *ggtranscript* R library (Gustavsson *et al*., 2022). Gene Ontology (GO) enrichment analyses were performed with the standalone version of *GeneMerge* v1.5 (Castillo-Davis & Hartl, 2003). The web application that deploys transcript expression profiles during de-etiolation was developed using the R Shiny web application framework.

### Alternative Splicing

To identify pairs of transcripts undergoing isoform switch between conditions, predict premature termination codons (PTCs) and Nonsense-Mediated mRNA Decay (NMD) sensitivity, we used IsoformSwitch (Vitting-Seerup *et al*., 2019). For isoform switch predictions a significance cutoff of 0.05 on the FDR corrected p-values and a difference in isoform fraction cutoff (DIFcuttof) of 0.1 were applied. TranSuite with default parameters was utilized to identify sequences of transcripts encoding proteins (Entizne *et al*., 2020).

### Validation of RNA-Seq profiles by qPCR

Total RNA was extracted from whole seedlings using the RNeasy Plant Kit (Qiagen). One microgram of total RNA was used as a template for reverse transcription with the iScript cDNA SynthesisKit (Bio-Rad). Before cDNA synthesis, genomic DNA in the samples was digested with DNase I (Thermo Fisher Scientific). One microliter of cDNA from each sample was used as template for real-time PCR analysis employing the primer combinations described in **S5 Data**. Reactions were performed with technical triplicates on CFX384 Real-Time System (Bio-Rad) quantitative PCR machine with iQ SYBR Green Supermix (Bio-Rad). To verify the purity of the PCR products, a melting curve was produced after each run. Arabidopsis genes, UBC (AT1G14400) and EF1 (AT1G07940), were used as reference genes and the relative transcript level was calculated by the ΔΔCT comparative quantification method (Livak and Schmittgen, 2001). for relative transcript quantification. Four biological replicates were used to calculate means and standard deviations of transcripts quantification.

### Protein Network Analysis

The structures of protein isoforms were predicted using AlphaFold2 v2.2.2 (Jumper *et al*., 2021), executed on NVIDIA V100 GPUs. To construct network models of protein isoforms, we used amino acid networks (Yan *et al*., 2014). VMD software (Humphrey *et al*., 1996) together with a custom Tcl script were employed to measure and average the distances between the alpha carbons of different amino acids within a cutoff of 7 Å for the five conformations predicted by AlfhaFold2 conformations. This resulted in weighted networks where each node represents an amino acid, and each edge represents an interaction between them. We applied the Infomap multilevel method (Rosvall & Bergstrom, 2008, 2011) to detect communities in the amino acid networks, with 10 trials for each network. We used Mapping Change and its online alluvial generator (Holmgren *et al*., 2023; Rosvall & Bergstrom, 2010) to analyze the changes in the structures of the protein isoforms.

### Molecular Dynamic analysis

To perform the molecular dynamics analysis between SIG5 protein isoforms and DNA, we used a DNA sequence of 75 nt that corresponds to a sequence located 930 bp upstream of the start codon of chloroplast encoded gene *PSBD* (ATCG00270), which includes −35-like and −10-like sequences and an AAG-box (Kanamaru *et al*., 2004) **(S1 File)**. We modelled the DNA conformation as B-DNA and the protein conformations were taken from the AlphaFold predictions. The orientations of the SIG5 protein isoforms with respect to the DNA were similar with respect to the main DNA axis. The protein and DNA molecules were initially separated by ~15 Å in distance. Parameters for atomic interactions were obtained from the CHARMM-36 force field (Best *et al*., 2012). An initial simulation for the protein-DNA system was performed during 8 ns where the Generalized Born model with a simple switching (GBSW) (Wang *et al*., 2022) as implemented in the CHARMM package (version 45b1) (Brooks *et al*., 2009). A cutoff of 17 Å was employed to compute electrostatic interactions explicitly that were switched off starting from 16 Å with a switching function as implemented in CHARMM. Then, the final structure of the implicit solvent simulation was employed to obtain a more realistic model where we included explicit TIP3P water molecules and salt at 150 mM, which consisted of Na^+^ and Cl^-^ ions. This step was performed with the GROMACS software (version 2023.1) (Páll *et al*., 2015, Pronk *et al*., 2013). The system was minimized by using harmonic restraints between 240 and 10 kcal/mol nm^-2^ for the backbone and side chain atoms. This step was followed by an equilibration period of 1 ns in the NVT ensemble with the Nosé-Hoover as a thermostat using a coupling constant of 1 ps. Finally, data was collected during 30 ns in the NPT ensemble where Parrinello-Rahman (Nosé & Klein., 1983) served as a barostat with a compressibility of 4.51 × 10^−5^ bar^−1^ and a coupling constant of 5 ps. Hydrogen bonds were constraint with LINCS algorithm (Hess *et al*., 1998). The electrostatic term of the potential energy function was computed by means of the particle mesh Ewald (PME) method (Essmann *et al*., 1995) with a grid size of 1.2 Å. VdW interactions were truncated at a 12 Å cutoff. All simulations were run at 303.15 K.

## Data Availability

Sequencing data from Hernández-Verdeja *et al*., (2022) are available at the European Nucleotide Archive (ENA) under accession number PRJEB47885.

The transcripts expression profiles during de-etiolation are available through an interactive web application at https://deetiolation-transcripts.serve.scilifelab.se/app/deetiolation-transcripts

## Acknowledgements

We thank the High Performance Computing Center North (HPC2N) at Umeå University for providing computational resources and valuable support during test and performance runs, and the UPSC Bioinformatics Facility for their computational infrastructure and dedicated assistance throughout our analyses.

## Supporting Information

**S1 Fig.**
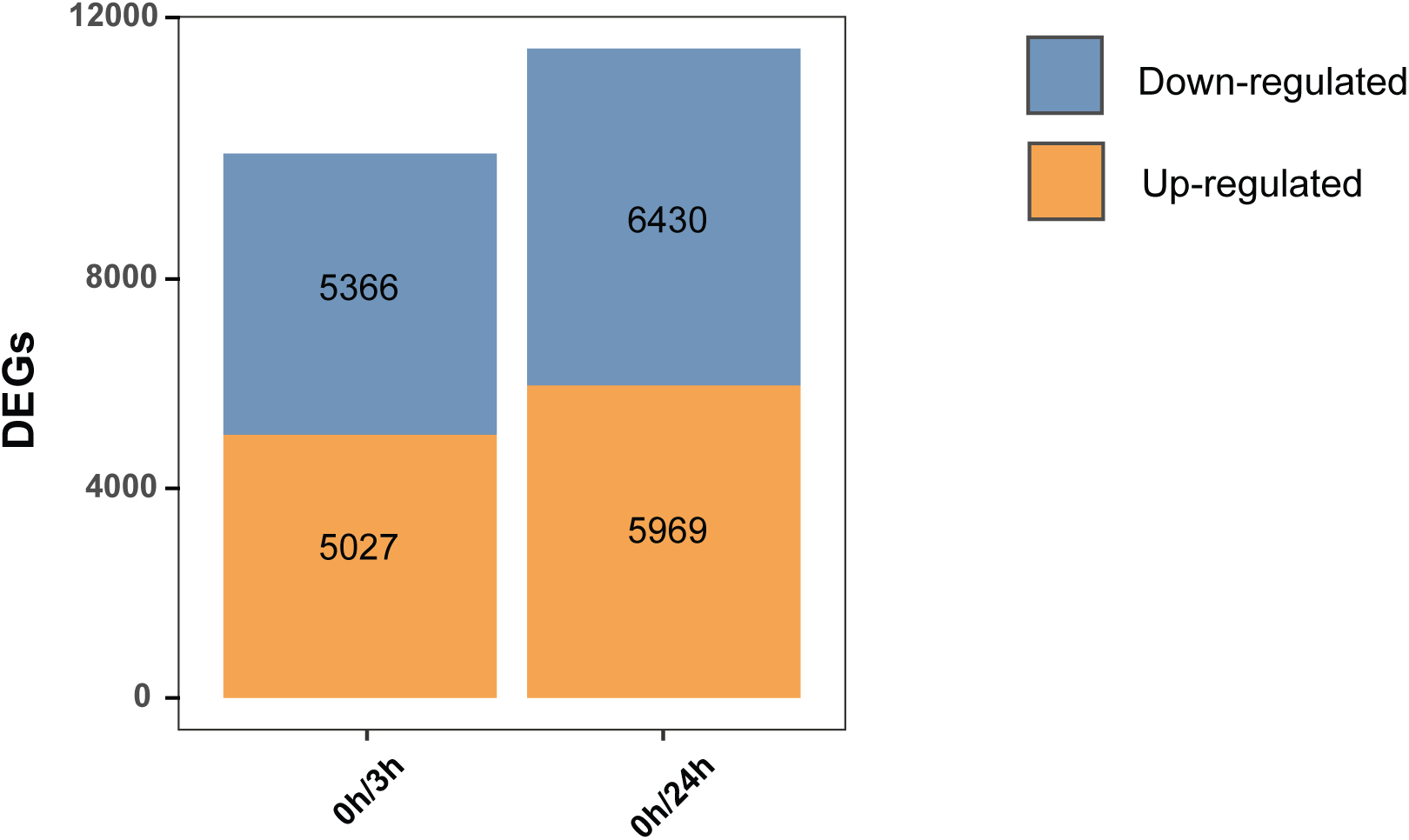
Differentially expressed genes in wild-type plants upon light exposure. Differentially expressed genes (DEGs) obtained by pairwise comparison. Up-regulated and Down-regulated DEGs (Adjusted Pvalue 0.05) are represented in orange and blue, respectively. Full lists of DEGs annotations, Log2FC and Adjusted P values for T0/3h and T0/24h comparisons are available in S6 and S7 Data respectively.

**S2 Fig.**
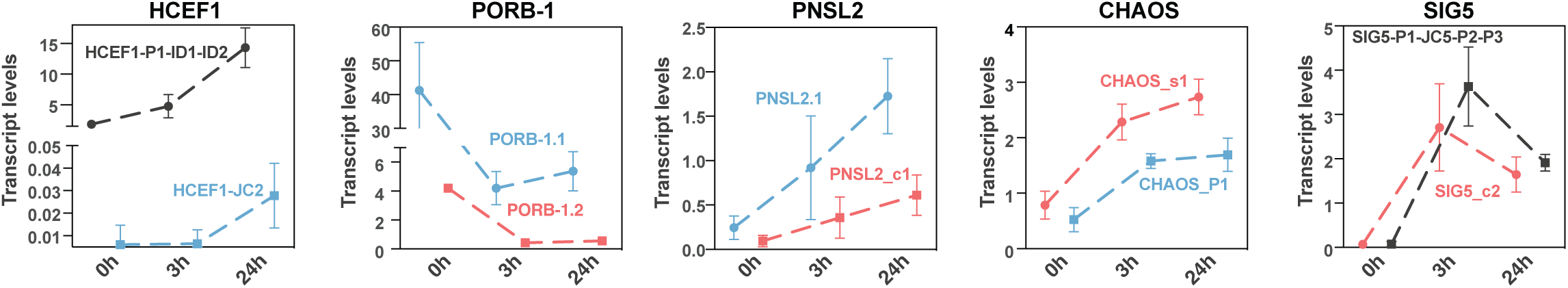
Validation of RNA-Seq profiles of photosynthesis related genes by qPCR. Each plot contains transcripts of a gene identified as undergoing isoform switch. Errors bars represent SD values. Names of the respective transcripts are shown in the same color of the profile that they represent. Black profiles represent cases where primers did not distinguish some transcripts, indicating the expression of more than one transcript. All associated transcripts are represented by the respective black profile.

**Figure S3.**
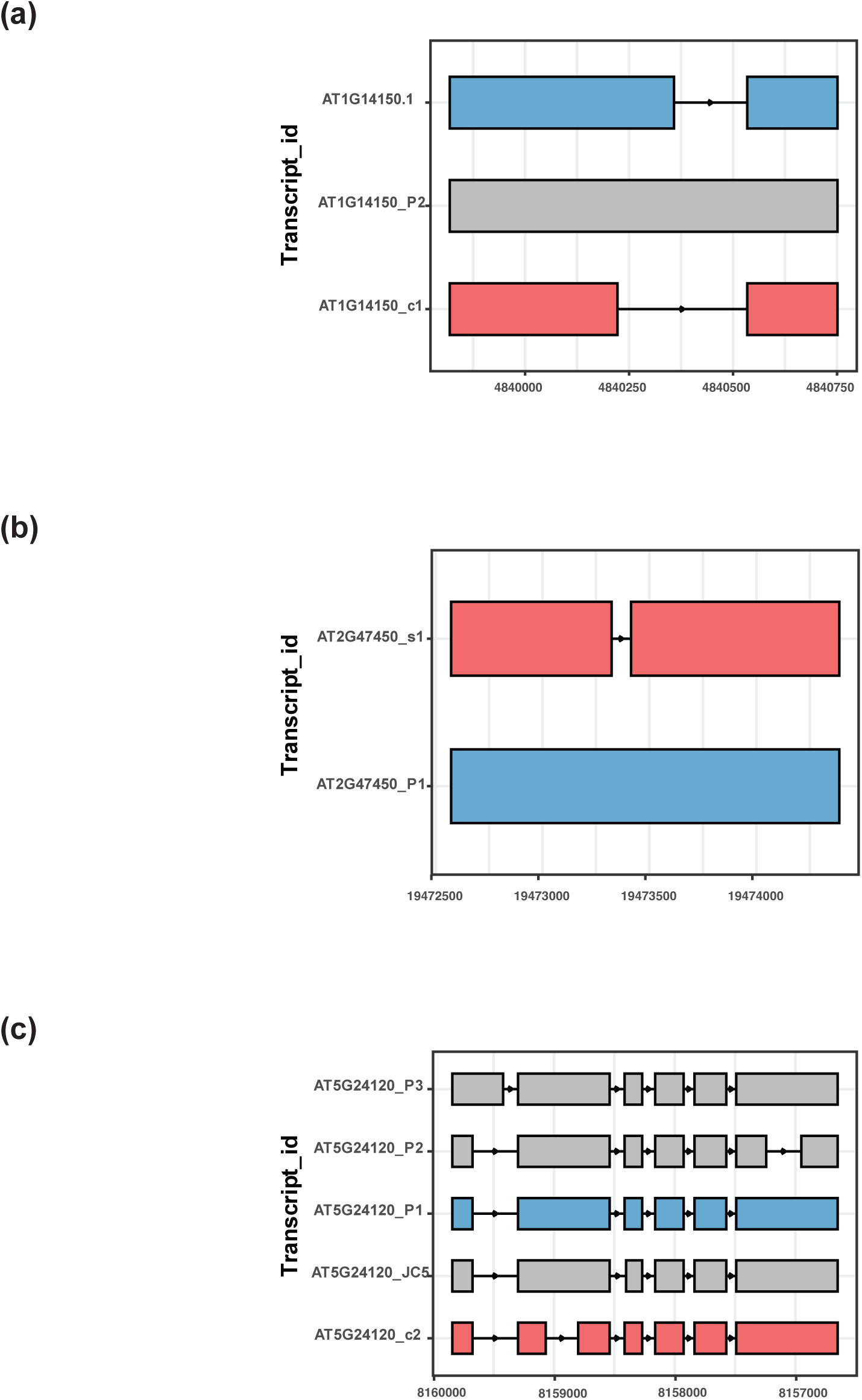
Transcript isoforms. Splice variants models for PNSL2, CHAOS and SIG5 (a, b,c, respectively). Total transcript models according to the AtRTD2 reference database (Zhang et al., 2017). Transcript models undergoing isoform switch with light exposure are represented in red and green. Transcripts that do not undergo isoform switch are represented in gray. Arrows indicate directionality of transcripts, from N terminal to the C terminal in the corresponding proteins.

**S4 Fig.**
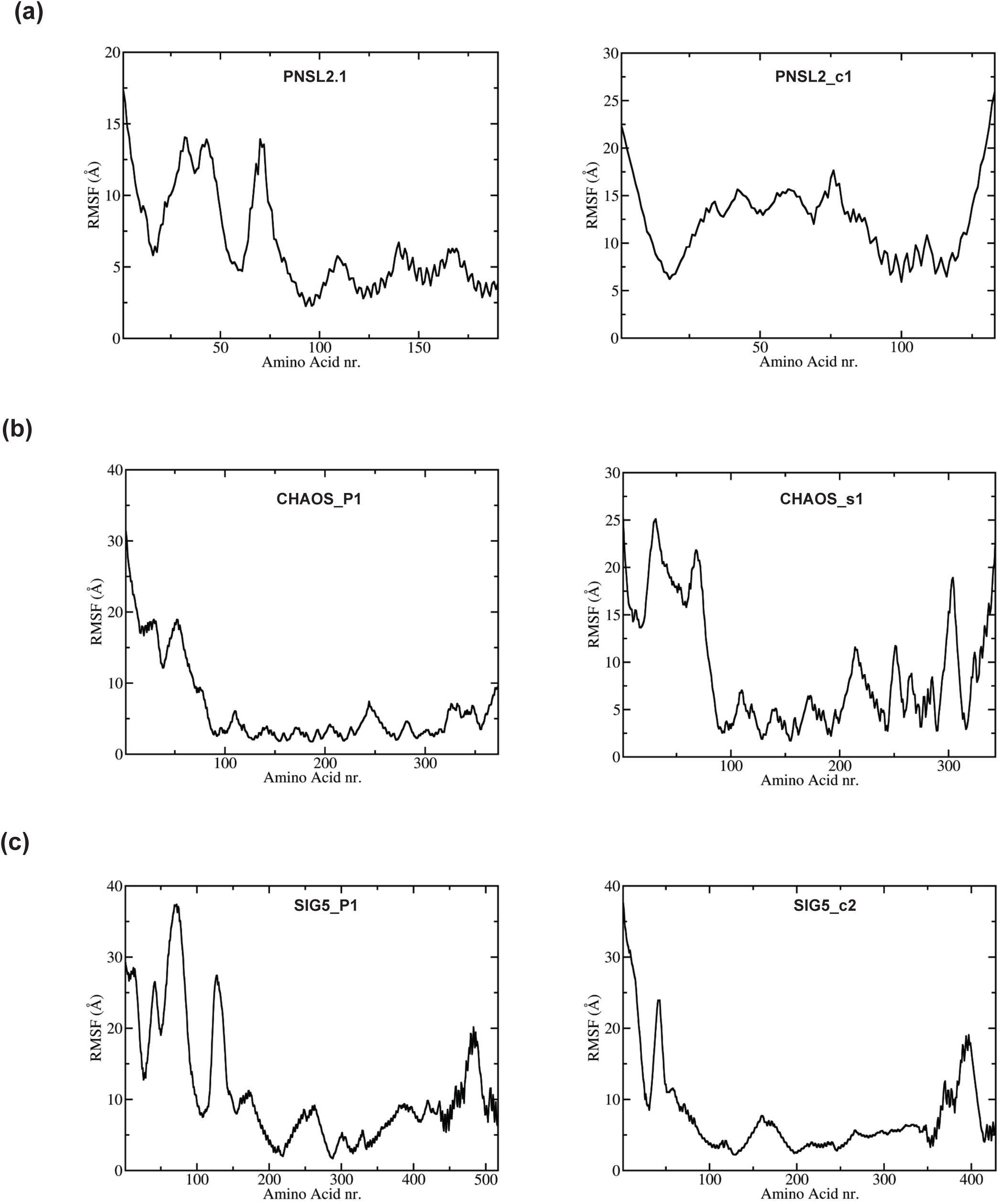
Dynamic flexibility profiles of protein isoforms. Representative AlphaFold protein structures were analyzed by Residue-wise Root Mean Square Fluctuation (RMSF) to represent the dynamic flexibility of each amino acid residue, providing information about protein isoforms local structural fluctuations. High RMSF values indicates regions of high flexibility. Lower values represent more stable and structured regions. (a) PNSL2, (b) CHAOS and (c) SIG5.

**S5 Fig.**
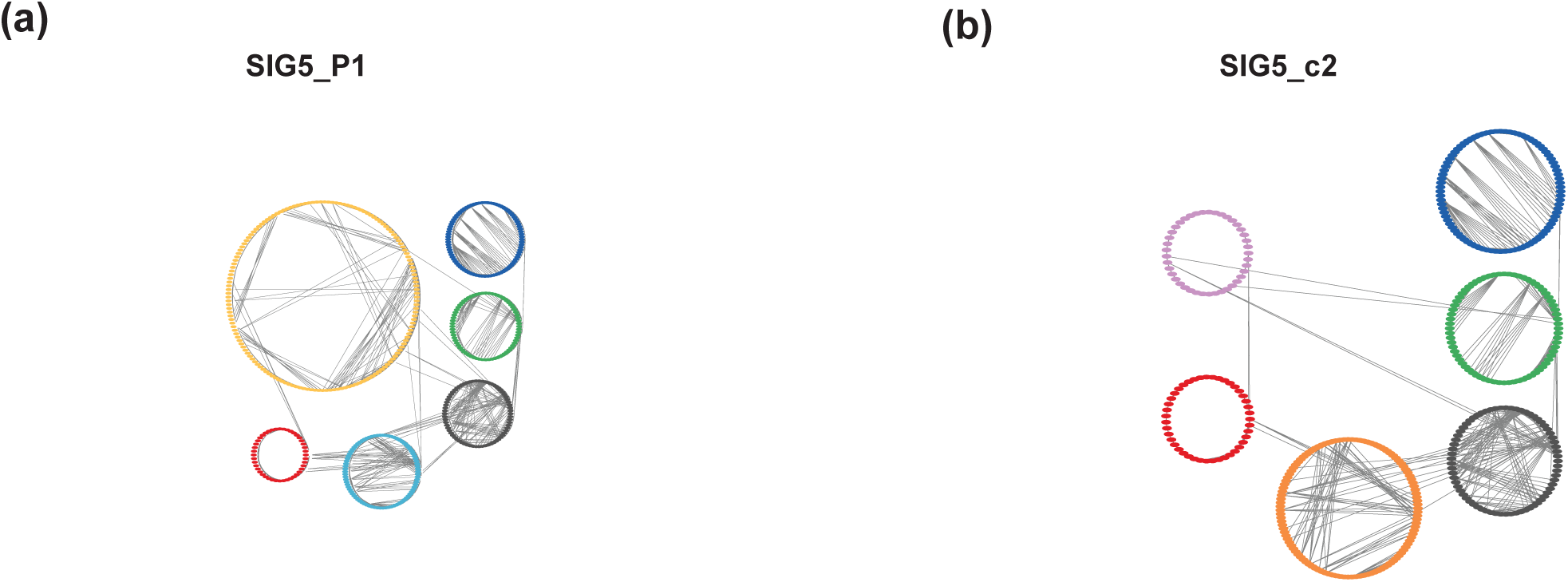
Network visualizations of SIG5 protein isoforms undergoing an isoform switch. Protein structure network representations of (a) SIG5_P1 and (b) SIG5_c2. Circles represent clusters that correspond to Infomap communities (see methods).

**S1 File.**
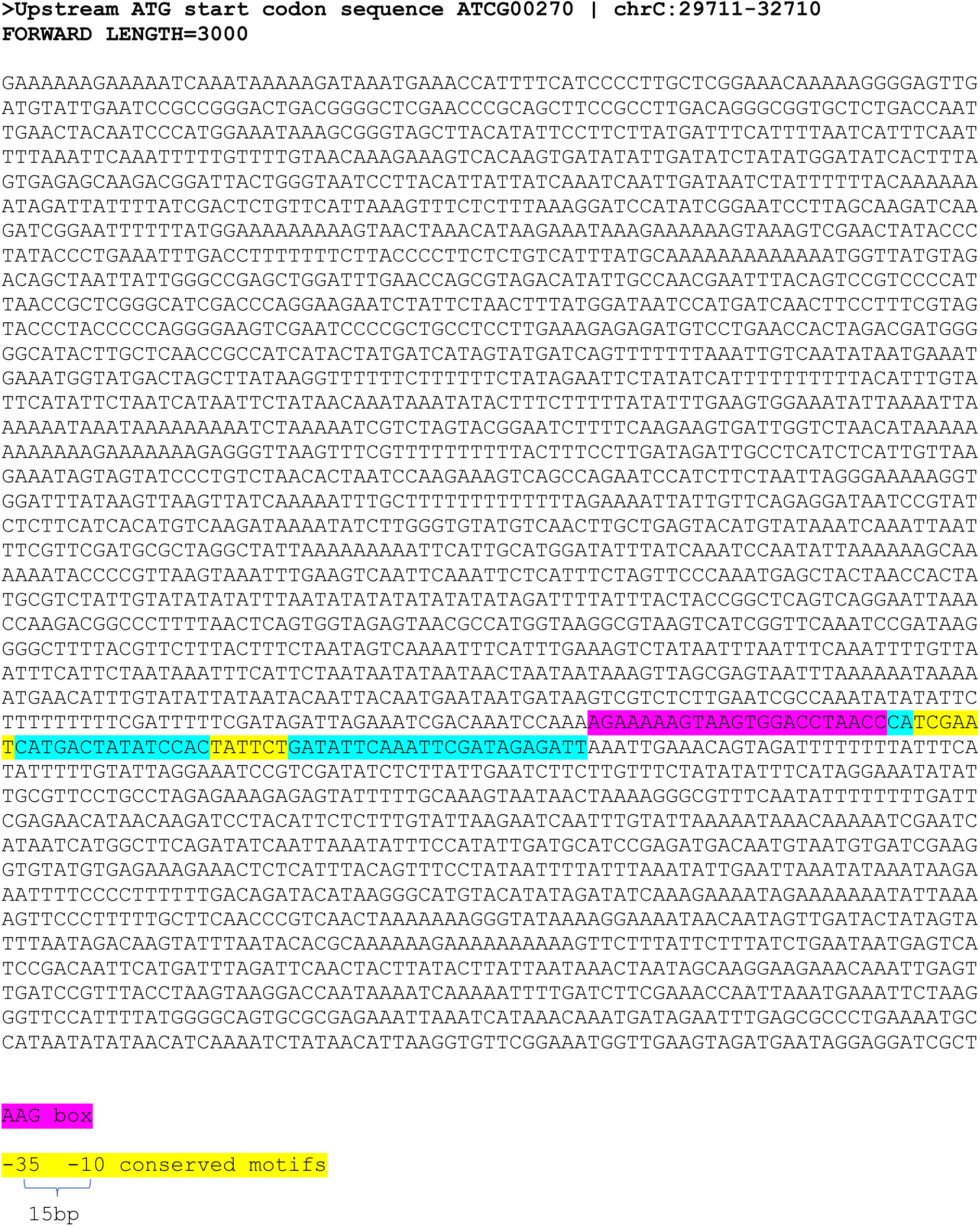
Upstream regulatory region of the chloroplast *psbD* gene (ATCG00270).

### Supplementary Data

**S1 Data.** Excel spreadsheet containing a list of differentially spliced genes (DSGs) in the wild type during de-etiolation, including annotations, alternative splicing mechanisms, the corresponding transcript isoforms involved, and the respective timepoints for comparison where changes were observed.

**S2 Data.** Excel spreadsheet containing a comprehensive list of DSGs changing between wild type and *gun1* during de-etiolation, including annotations, alternative splicing mechanisms, the corresponding transcript isoforms involved, and the respective timepoints for comparison where changes were observed.

**S3 Data.** Excel spreadsheet containing 40 genes that were identified. All these genes are genes related to chloroplast or photosynthesis that exhibit transcript changes in wild-type plants during de-etiolation, and exhibit transcript alterations in *gun1-102* plants compared to the wild-type.

**S4 Data.** Excel spreadsheet containing 40 identified genes and their associated transcripts that undergo isoform switch during de-etiolation. Transcripts with predicted premature termination codons (PTCs) are indicated.

**S5 Data.** Excel spreadsheet containing primer sequences used for validation of RNA-Seq profiles by qPCR.

**S6 Data.** Excel spreadsheet containing differentially expressed genes in wild-type plants upon light exposure obtained by comparing 0h versus 3h. DEGs annotations, Log2FC and Adjusted P values are available.

**S7 Data.** Excel spreadsheet containing differentially expressed genes in wild-type plants upon light exposure obtained by comparing 0h versus 24h. DEGs annotations, Log2FC and Adjusted P values are available.

## Notes

### Competing Interest Statement

The authors have declared no competing interest.

